# Restoring memory by optogenetic synchronization of hippocampal oscillations in an Alzheimer’s disease mouse model

**DOI:** 10.1101/363820

**Authors:** Eleonora Ambrad Giovannetti, Stefanie Poll, Daniel Justus, Hiroshi Kaneko, Falko Fuhrmann, Julia Steffen, Stefan Remy, Martin Fuhrmann

**Affiliations:** Neuroimmunology and Imaging Group, German Center for Neurodegenerative Diseases (DZNE); Neuronal Networks Group, German Center for Neurodegenerative Diseases (DZNE), Bonn, Germany; Department of Epileptology, University of Bonn Medical Center, Bonn, Germany

## Abstract

Disrupted neural oscillations are a feature of Alzheimer’s disease (AD). We observed reduced frequency of theta oscillations in the hippocampal local field potential (LFP) in a mouse model of beta-amyloidosis. By restoring the temporal organization of theta oscillations using LFP-guided closed-loop optogenetic stimulation of parvalbumin-positive interneurons, we could rescue memory deficits of APP/PS1 mice in the novel object recognition test.

---

Aberrant neuronal theta and gamma oscillations and neuronal hyperexcitability have been observed in patients and mouse models of Alzheimer’s disease (AD)^1–8^. Both pathophysiological phenomena may have detrimental effects on cognitive function. While aberrant oscillatory patterns are expected to interfere with mnemonic processing by disrupting the temporal coding within neuronal ensembles^9^, neuronal hyperexcitability is likely resulting in pathophysiological alterations of the neuronal input to output transformation^6, 10^. Novel interventional strategies should therefore act effectively at the network and the single cell level^11^. During mnemonic processing within the entorhinal-hippocampal circuitry, theta oscillations in the 4-12 Hz frequency range are prominent in rodents and in humans^9^. They are driven by rhythmic activity of septo-hippocampal projections, which target hippocampal interneurons^9^. A reduced theta frequency has been observed in AD mouse models and patients^12–14^. Notably, we and others have previously shown that AD-like pathology results in inhibitory interneuron dysfunction^15^. Specifically, parvalbumin positive (PV^+^) interneurons were critically affected in humans and mouse models of AD^7, 16, 17^. Increasing their inhibitory function rescued spatial memory deficits^7^ and reduced Aβ-plaque load in the visual cortex^3^. Increasing neuronal inhibition was also beneficial in human MCI patients and mouse models following administration of levetiracetam, a pro-inhibitory antiepileptic drug^18, 19^. Mechanistically, increasing inhibitory function of specific interneurons may represent a way to counteract both aberrant network activity patterns and increased cellular excitability. Therefore, we designed a closed-loop theta detection and optogenetic stimulation approach based on continuous monitoring of the hippocampal local field potential. This approach was targeted specifically at PV^+^ interneurons and resulted in a simultaneous shift of theta frequency towards more physiological levels and an increase in hippocampal PV^+^ mediated inhibition. We hypothesized that such an approach could be effective in improving memory in an APPswe/PS1∆E9 (APP/PS1) mouse model of Alzheimer’s disease – restoring the temporal organization of theta oscillations by shifting them to a frequency range that is observed under physiological conditions.

First, to quantify differences in hippocampal oscillations between wild type and APP/PS1 transgenic animals, we recorded CA1 local field potential in the bilateral str. radiata during open field exploration (Figure 1a, b). In the transgenic group, the peak theta frequency during locomotion was significantly decreased by 0.4 Hz compared to wild type animals (Figure 1c; *WT*: 8.3 ± 0.07 Hz, n = 24 mice; *APP/PS1*: 7.9 ± 0.07 Hz, n = 21 mice). Neither the mean power of theta oscillations nor their mean duration was significantly altered between the two groups (Supplementary fig. 1). The mean running speed was significantly higher in transgenic animals (Figure 1d; *WT*: 6.3 ± 0.40 cm/sec, n=24 mice; *APP/PS1*: 7.9 ± 0.44 cm/sec). Nevertheless, although theta frequency is well known to positively correlate with running speed^20^, the peak theta frequency was reduced. Travelled distance and time spent in the center of the arena were not different between the groups (Figure 1e, f).

**Figure 1.**
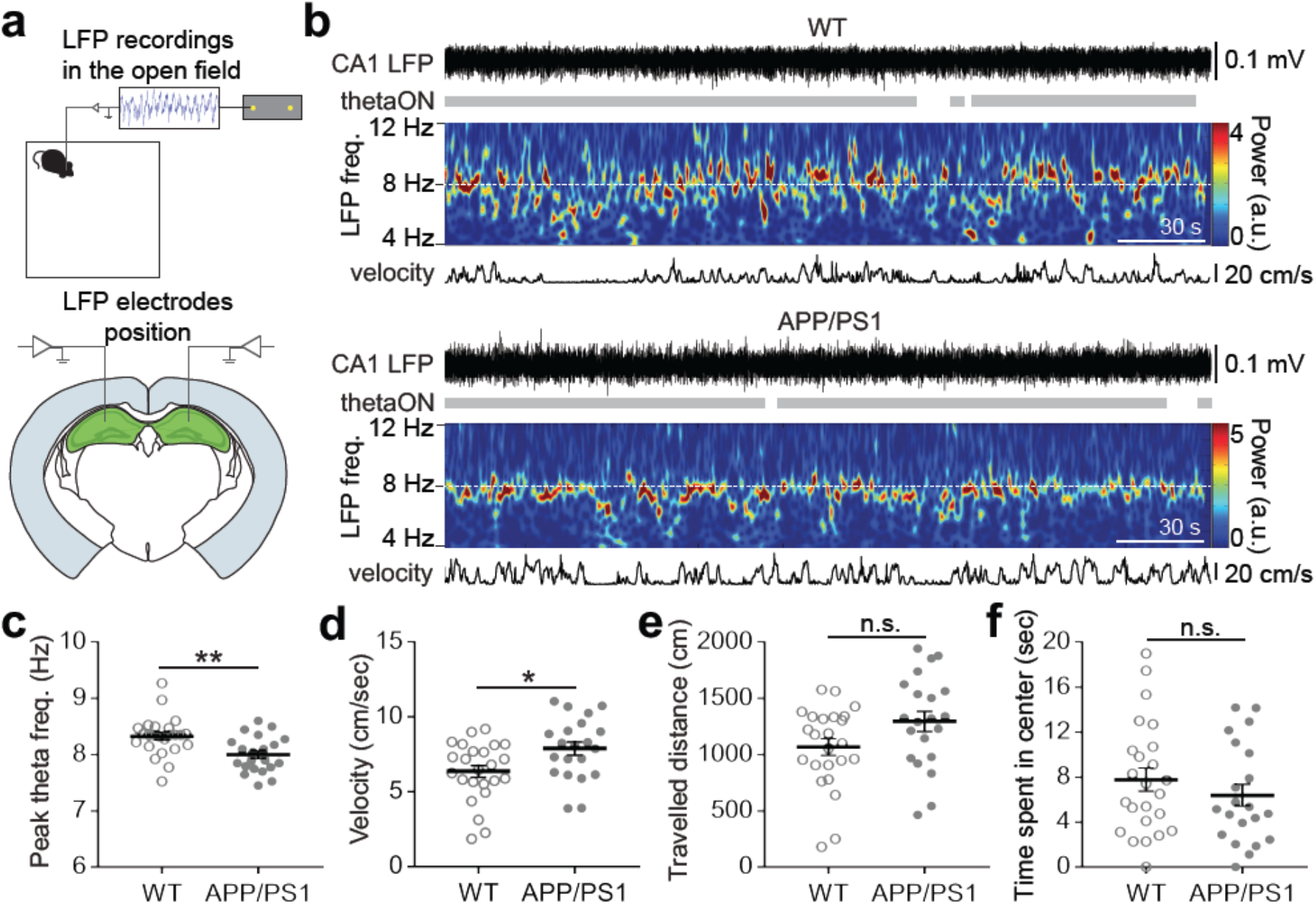
Slower theta oscillations in aged APP/PS1 mice. **(a)** Schematics of LFP recording configuration. **(b)** Exemplary spectrograms showing the slower peak theta frequency recorded in APP/PS1 compared to wild type (WT) mice during open field exploration. Theta ON refers to the intervals of increased theta power detected during offline analyses. **(c)** Peak theta frequency of WT and APP/PS1 mice,detected during ThetaON intervals (^**^p<0.01; unpaired Student’s t-test). **(d)** Mean velocity during epochs of locomotion (point velocity >0.5 cm/sec); ^*^p<0.05, unpaired Student’s t-test. **(e)** Total travelled distance. **(f)** Time spent in the center of the arena. For **e** and **f**, n.s.= p>0.05, not significant; unpaired Student’s t-test. For **c** to **f**, WT: n= 24, APP/PS1: n=21. Dots represent individual mice, bars show mean ± SEM. Individual mouse values result from the average of 2-6 recordings of 3 min each.

Next, we tested the effect of optogenetic stimulation of PV^+^ interneurons on the hippocampal LFP. Therefore, we bilaterally injected *rAAV2-Ef1a-DIO C1V1 (t/t)-TS-mCherry* into the CA1 subregions of PV-Cre mice on wild type or APP/PS1 genetic background, thereby restricting the expression of C1V1 to PV^+^ interneurons (Figure 2a, b). As controls, an additional group of mice with the identical genetic background was injected with a construct encoding a loxP-flanked fluorescent reporter protein (*rAAV1-CAG-FLEX-tdTomato*). Bilateral optical fibers were implanted, terminating above the transfected region (Figure 2a, b; Supplementary figure 5). Light stimulation significantly increased the power of the hippocampal oscillations at the frequency bands at which stimulation was performed in C1V1 injected animals, but not in the Sham injected group (Figure. 2c and 2d; *Sham*: n = 4 to 5 animals; *C1V1*: n = 6 animals; Supplementary figure 4).

**Figure 2.**
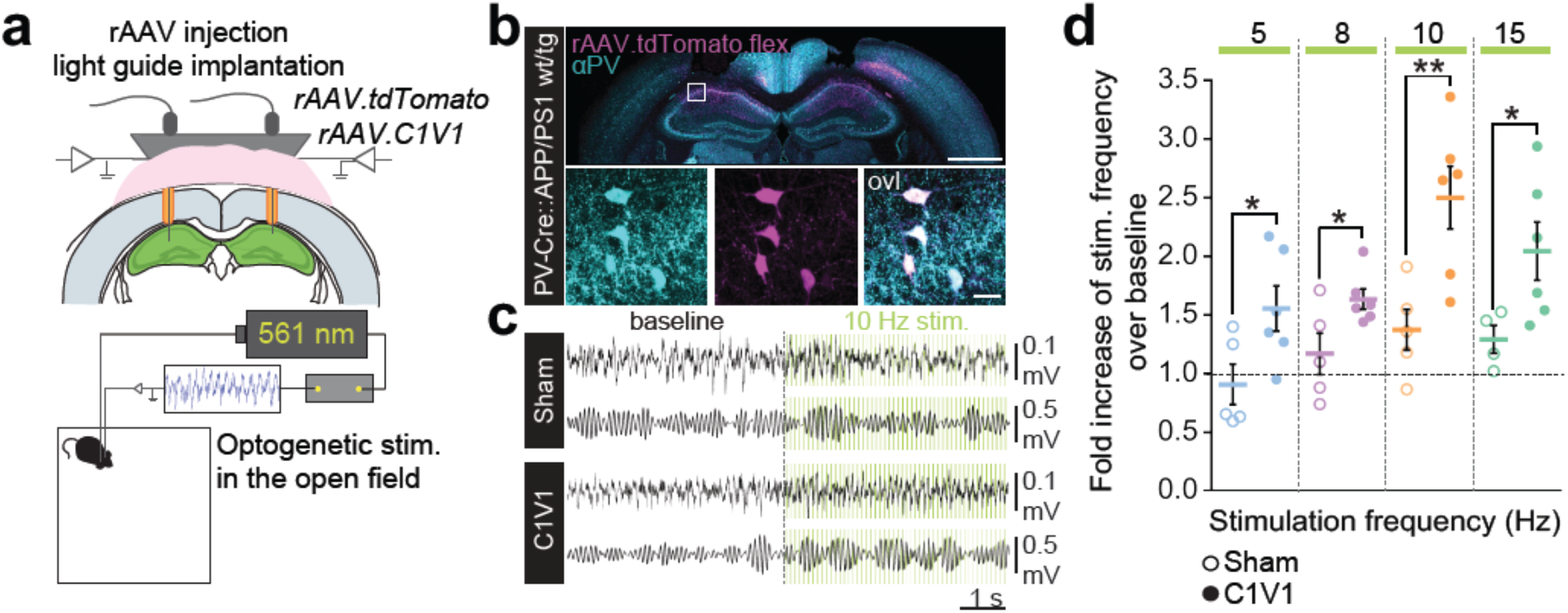
Optogenetic stimulation of PV^+^ interneurons drives LFP frequencies. **(a)** Schematics of experimental setup. **(b)** Exemplary picture of rAAV expression in PV^+^ interneurons in the hippocampus. Scale bars: 1 mm (upper panel) and 20 μm (lower panel), ovl = overlap **(c)** Exemplary LFP traces recorded from mice injected with rAAV-tdTomato (Sham) and rAAV-C1V1 (C1V1) during a 10 Hz light (561 nm) stimulation protocol. Upper traces: filtered LFP, 3-200 Hz. Lower traces: filtered LFP, 8-12 Hz. **(d)** Fold increase of stimulation frequency over baseline (Sham: n= 5 mice for all frequencies but 15 Hz, where n=4, C1V1: n= 6 mice for all frequencies; ^*^p<0.05, ^**^p<0.01; unpaired Student’s t-tests). The dashed line represents the baseline to which the fold increase was normalized. Dots represent individual mouse values, bars show mean ± SEM. Individual mouse values result from the average of 4-6 stimulation protocols.

There are two plausible and mutually non-exclusive mechanisms by which optogenetic stimulation of PV^+^ interneurons could lead to an improvement of memory performance in APP/PS1 mice: First, the stimulation of PV^+^ interneurons in the theta range is expected to lead to a rhythmic recruitment of these interneurons. As demonstrated before, such a rhythmic, synchronous firing of many PV^+^ interneurons will likely result in a stimulation-frequency dependent field potential oscillation^20, 21^. It is possible that by (i) shifting the mean theta frequency towards more physiological levels in APP/PS1 mice and by (ii) increasing the rhythmicity of the oscillation, the temporal coding of memory-relevant information would be improved. The relative timing of synaptic input and action potential firing in relation to the phase of the theta oscillation has been implicated to be relevant for the encoding and retrieval of memories (for ref. see^22, 23^). Thus, by defining the time window at which inhibition permits synaptic input to cause action potential firing, the effectiveness of encoding could be enhanced. Second, neuronal hyperexcitability in AD models has been shown to profoundly alter the input-output conversion of neurons resulting in higher firing rates and a shift of the output pattern towards more burst firing^6, 10^. Since the synchronous and repetitive stimulation of interneurons is expected to result in a sustained increase in inhibition during the stimulation periods, it is expected to cause a shift of the input-output function in hippocampal neurons towards a reduced spike output at a given input strength^24^. We therefore designed an experimental approach that aimed at distinguishing the effects of increased rhythmicity of pyramidal cell output on memory performance from effects caused by a general increase in inhibitory strength. Our approach aimed at maintaining an equal inhibitory strength in two dissimilar stimulation patterns: Both stimulation patterns consisted of equal numbers of stimuli, but differed in the temporal precision at which pulses were delivered. In the “rhythmic” stimulation protocol the interpulse interval was kept stable so that the light pulses were equally spaced. The “arrhythmic stimulation” protocol introduced a random jitter in the interpulse interval duration that did not exceed 30 ms. Before applying these protocols in a closed-loop optogenetic feedback stimulation *in vivo*, we carried out whole-cell patch-clamp recordings of PV^+^ and CA1 pyramidal neurons (PYR) in acute slices of wild type PV-Cre mice injected with *rAAV2-Ef1a-DIO C1V1 (t/t)-TS-mCherry* (Figure 3a). These experiments confirmed a) the high reliability of light-induced action potential generation in PV+ interneurons, b) an inhibitory effect of PV^+^ stimulation on the membrane potential of CA1 pyramidal neurons and most importantly c) no difference in the strength of inhibition evoked by both stimulation protocols (Figure 3b-d). The latter was determined by a comparison of IPSC amplitudes evoked by repetitive stimulation. Subsequently, these protocols were implemented into closed-loop optogenetic feedback stimulation paradigms; two groups of mice were subjected to either the “rhythmic” or the “arrhythmic” feedback stimulation during a novel object recognition test (NOR). In both cases, the optogenetic feedback stimulation was driven by the endogenous theta oscillations recorded in the LFP of the behaving mice. In particular, an increase in the theta oscillation power would trigger the optogenetic stimulation at a frequency equal to the one detected in the LFP, plus an additional 0.5 Hz to compensate for the deficit measured in APP/PS1 transgenic mice (Figure 3e). The NOR was carried out 24 h after the memory acquisition phase (Figure 3f), and discrimination indices (DI) were calculated to describe the degree of memory retention (*See Methods*). The rhythmic stimulation protocol resulted in an improved recognition memory in C1V1-injected PV-Cre::APP/PS1 mice in contrast to Sham-injected PV-Cre::APP/PS1 mice (Figure 3g; *APP/PS1-Sham*: DI = 41.8% ± 8.18%; n = 11 mice; *APP/PS1-C1V1*: DI = 66.6 ± 7.46%, n = 11 mice). Interestingly, the rhythmic optogenetic stimulation of PV^+^ interneurons in the C1V1-injected wild type group impaired recognition memory compared to Sham-injected controls (Figure 3g; *WT-Sham*: DI = 73.2% ± 7.03%, n = 8, *WT-C1V1*: DI = 39.8% ± 6.18%, n = 11). Notably, while the rhythmic optogenetic stimulation could rescue impaired recognition memory, the arrhythmic stimulation protocol was not sufficient to improve novel object discrimination of C1V1-injected PV-Cre::APP/PS1 mice compared to Sham-injected PV-Cre::APP/PS1 mice (Fig. 3h; *APP/PS1-Sham*: DI = 46.3% ± 14.3, n = 7; *APP/PS1-C1V1*: DI = 49.4% ± 13.4%, n = 5). Mean travelled distance was comparable between all experimental groups, indicating no effect of the light stimulation on locomotor activity (Supplementary figure 2). Light delivery *per se* had no effect on the LFP of Sham-injected mice subjected to the rhythmic feedback stimulation (Supplementary figure 3 a-g), nor on the LFP of C1V1 injected mice subject to the arrhythmic feedback stimulation (Supplementary figure 3 h-k).

**Figure 3.**
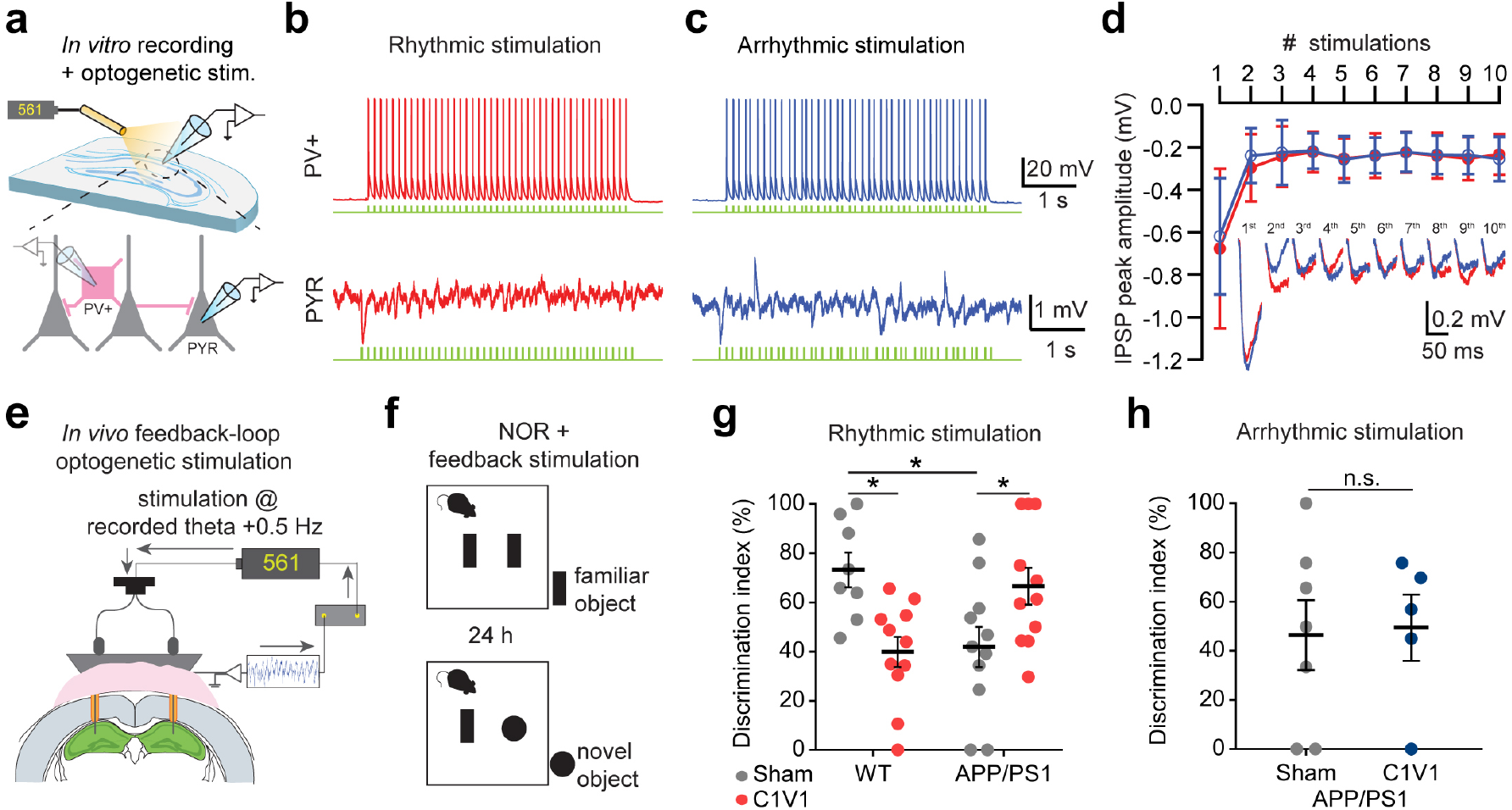
Enhanced neural synchronization via PV^+^ interneuron stimulation improves memory. **(a)** Schematics of the *in vitro* recording and optogenetic stimulation configuration. **(b, c)** Exemplary recordings from a C1V1-expressing PV+ interneuron and a CA1 pyramidal neuron (PYR) during rhythmic (b) and arrhythmic (c) 8.5 Hz light stimulation. **(d)** Average peak IPSP amplitudes recorded from CA1 pyramidal neurons plotted against the number of optogenetic stimulations comparing rhythmic (red) and arrhythmic (blue) stimulation of C1V1-expressing PV+ interneurons. Inset: individual example. **(e)** Schematics illustrating the *in vivo* recording and feedback optogenetic stimulation configuration. **(f)** Schematics of the novel object recognition (NOR) test with a 24h interval time. Optogenetic feedback stimulation was carried out upon theta frequency detection during the NOR test. **(g, h)** Discrimination index of wild type (WT) and APP/PS1 mice expressing either C1V1 or tdTomato (Sham) in PV^+^ interneurons. Mice performed the NOR task during rhythmic (g) or arrhythmic (h) feedback-loop optogenetic stimulation of PV+ interneurons; ^*^p<0.05, two-way ANOVA with Holm-Sidak’s correction (Interaction factor= 29.3% of total variation, ^***^p<0.001; row and column factors= 0.18% and 0.64% of total variation, p>0.05); n.s.= p>0.05, not significant; unpaired Student’s t-test. (For **g**, WT-Sham: n= 8, APP/PS1-Sham: n= 11, WT-C1V1: n= 11, APP/PS1-C1V1: n= 11. For **h**, APP/PS1-Sham: n=7, APP/PS1-C1V1: n=5. Dots represent individual mouse values, bars show mean ±SEM).

Taken together, we found that the peak frequency of theta oscillations in APP/PS1 mice was significantly reduced when compared to wild type littermates (albeit their higher movement speeds), a finding which is consistent with previous reports in patients and AD mouse models^12–14^. Most remarkably, the closed-loop rhythmic optogenetic stimulation of hippocampal PV^+^ interneurons effectively rescued the impaired recognition memory of APP/PS1 mice during the novel object recognition task bringing it back to levels comparable to wild type mice. This rescue was not observed upon arrhythmic stimulation, suggesting that memory processes may benefit from an optogenetic enhancement of theta rhythmicity and an associated improvement in temporal coding rather than from an increase in bulk inhibition alone. The result that optogenetic treatment of wild type mice had detrimental effects on memory performance underscores that, physiologically, the temporal coding of neuronal ensemble activation is optimally tuned, in contrast to pathophysiological conditions. Thus, it is possible that shifting the frequency of the endogenous theta rhythm towards higher frequencies and increasing bulk inhibition brought this optimized tuning in disarray, while the same procedure effectively restored the prevalent temporal asynchrony and hyperexcitability of the hippocampal circuitry, favoring more effective memory processing in APP/PS1 mice.

In conclusion, we showed that restoring the disrupted theta rhythmicity measured in PV-Cre::APP/PS1 transgenic mice by means of rhythmic optogenetic PV^+^ interneuron stimulation rescued long-term memory deficits in a novel object recognition task. Electrical deep brain stimulation has been used in several brain regions in patients and positive effects on memory performance have been reported in studies in humans including AD patients^25^. However, it is likely that the performance enhancing effect of deep brain stimulation is multi-factorial and mechanistically not precisely defined^25, 26^. Our results using optogenetic stimulation now identify PV^+^ interneurons as a target and a mean to restore neural synchronicity. Ultimately, this may serve to guide the development of more effective and specific interventional stimulation strategies in humans.

## Acknowledgements

This work was supported by the DZNE, grants from the Deutsche Forschungsgemeinschaft (SFB1089 B01, C01, B06, C06 to M.F. and S.R.), ERA-NET MicroSynDep, and the CoEN initiative (3018).

## Author contributions

E.A.G. performed *in vivo* experiments, analyzed the data and prepared figures. H.K. performed the patch-clamp recordings with optogenetic stimulation in acute brain slices and analyzed the data. S.P. and F.F. provided assistance for surgical procedures. J.S. provided technical assistance with perfusions and histology. D.J. and S.R. designed the closed-loop stimulation, D.J. wrote the custom IgorPro scripts for closed-loop detection and stimulation, and the MATLAB scripts for field potential analysis. S.R, M.F. and E.A.G. wrote the manuscript, with contributions from all co-authors. S.R. and M.F. coordinated research. M.F. supervised the research project and designed the experiments.

---

**Supplementary figure 1.**
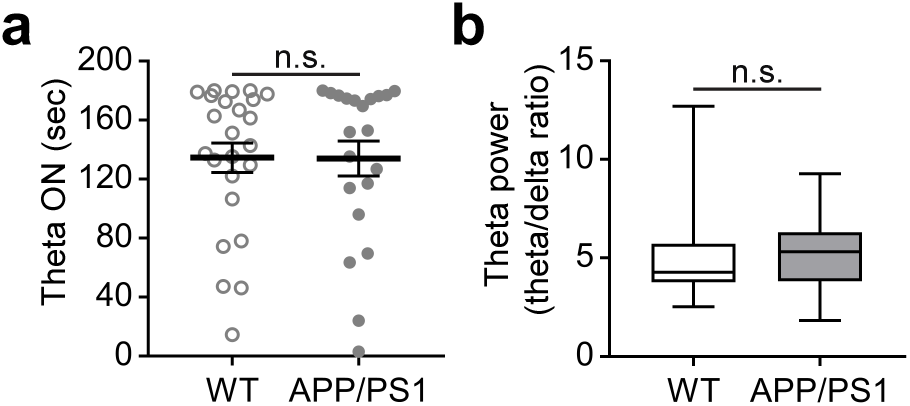
Duration and power of theta oscillations were not altered in APP/PS1 transgenic mice. **(a)** Average duration of theta oscillations during active exploration in the open field (2-6 recordings of 3 min per mouse); unpaired Student’s t-test. The graph represents individual mouse values, mean ± SEM. **(b)** Amplitude of the recorded theta oscillations defined as the ratio of theta band (5-12 Hz) over the delta band (4-5 Hz); Mann-Whitney test. Whisker plots represent median, 25^th^ and 75^th^ percentiles. For **a** and **b** WT: n=24, APP/PS1: n=21; n.s.= not significant.

**Supplementary figure 2.**
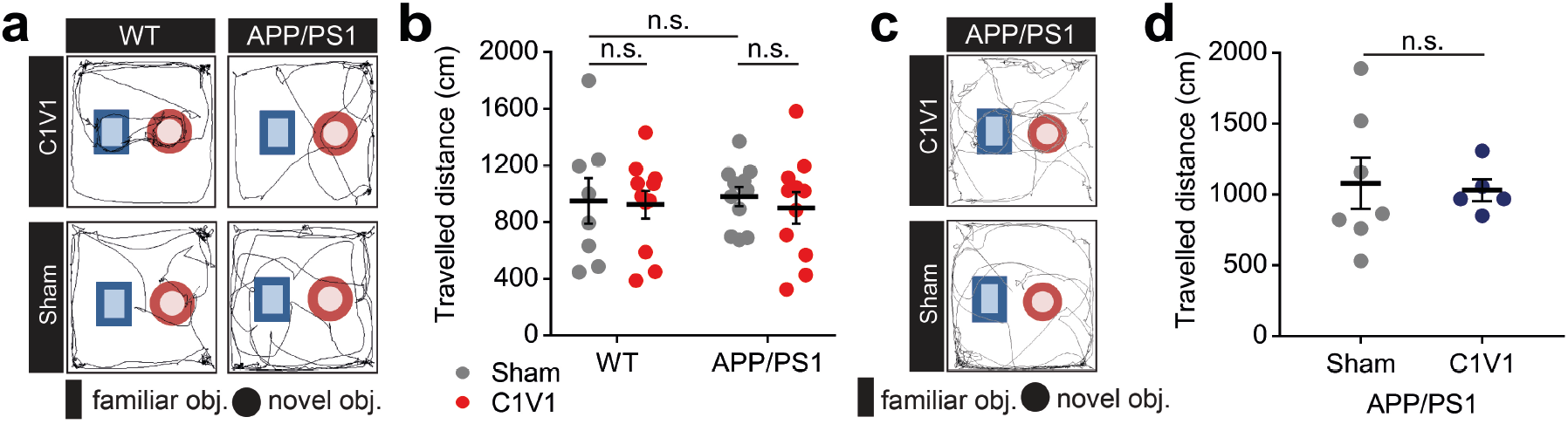
Travelled distance during the NOR test was unaltered upon feedback loop stimulation and unchanged between genotypes. **(a)** Rhythmic stimulation: Exemplary traces of single mice exploring novel (red circle) or familiar (blue rectangle) object in the open field. **(b)** Total travelled distance during the test; n.s.= not significant, two-way ANOVA with Holm-Sidak’s correction. **(c)** Arrhythmic stimulation: Exemplary traces of single mice exploring novel (red circle) or familiar (blue rectangle) object in the open field. **(d)** Total travelled distance during the test; n.s.= p>0.05, not significant; unpaired Student’s t-test. Dots represent individual mouse values, bars show mean ± SEM.

**Supplementary figure 3.**
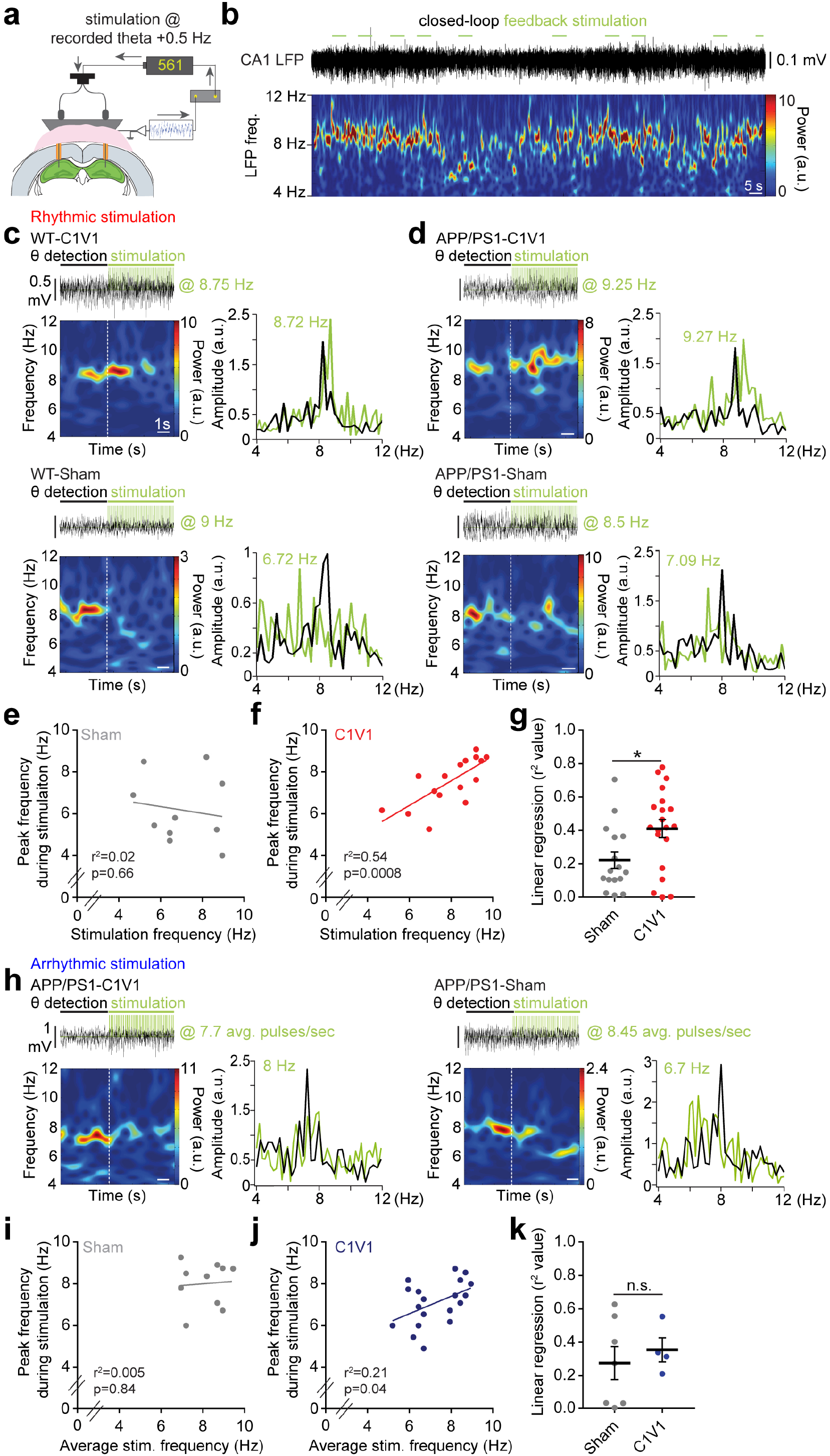
Effect of the closed-loop feedback stimulation of PV^+^ interneurons on the hippocampal LFPs.. **(a)** Schematics of the closed feedback-loop optogenetic stimulation. **(b)** Exemplary LFP trace and respective spectrogram showing the activation of the closed loop throughout one recording. **(c-d)** *Rhythmic stimulation*: Exemplary LFP epoch, respective spectrograms and fast Fourier transform of a single stimulation event driven by the endogenous theta frequency detected in the LFP of C1V1-(upper panel) and Sham-injected (lower panel) WT mice (c). Respective examples for APP/PS1 mice (d). Scale bars: 0.5 mV and 1 sec for all spectrograms. Note that in Sham-injected mice (WT-Sham) the stimulation has no effect on the LFP. **(e-f)** Exemplary x-y plot showing the stimulation frequency on the (x-axis) and the respective peak frequency (y-axis) recorded during stimulation epochs for a Sham-(e) and a C1V1-(f) injected mouse. **(g)** Cumulative r^2^ value for Sham- and C1V1-injected animals (Sham: n= 16; C1V1: n= 20 mice; ^*^p<0.05, unpaired Student’s t-test). **(h)** *Arrhythmic stimulation:* Exemplary LFP epoch, respective spectrograms and fast Fourier transform of a single stimulation event driven by the endogenous theta frequency detected in the LFP of APP/PS1 mice. Scale bars: 1 mV and 1 sec for all spectrograms. **(i-j)** Exemplary x-y plot showing the average stimulation frequency on the (x axis) and the respective peak frequency (y axis) recorded during arrhythmic stimulation epochs for a Sham (i) and a C1V1 (j) injected mouse. **(k)** Cumulative r^2^ value for Sham- and C1V1-injected animals (Sham: n= 7; C1V1: n= 4 mice; n.s.= p>0.05, not significant, unpaired Student’s t-test). For **e**-**f** and **i-j**: each dot represents a single stimulation event. For **g** and **k**: dots represent individual mouse values, bars show means ± SEM.

**Supplementary figure 4.**
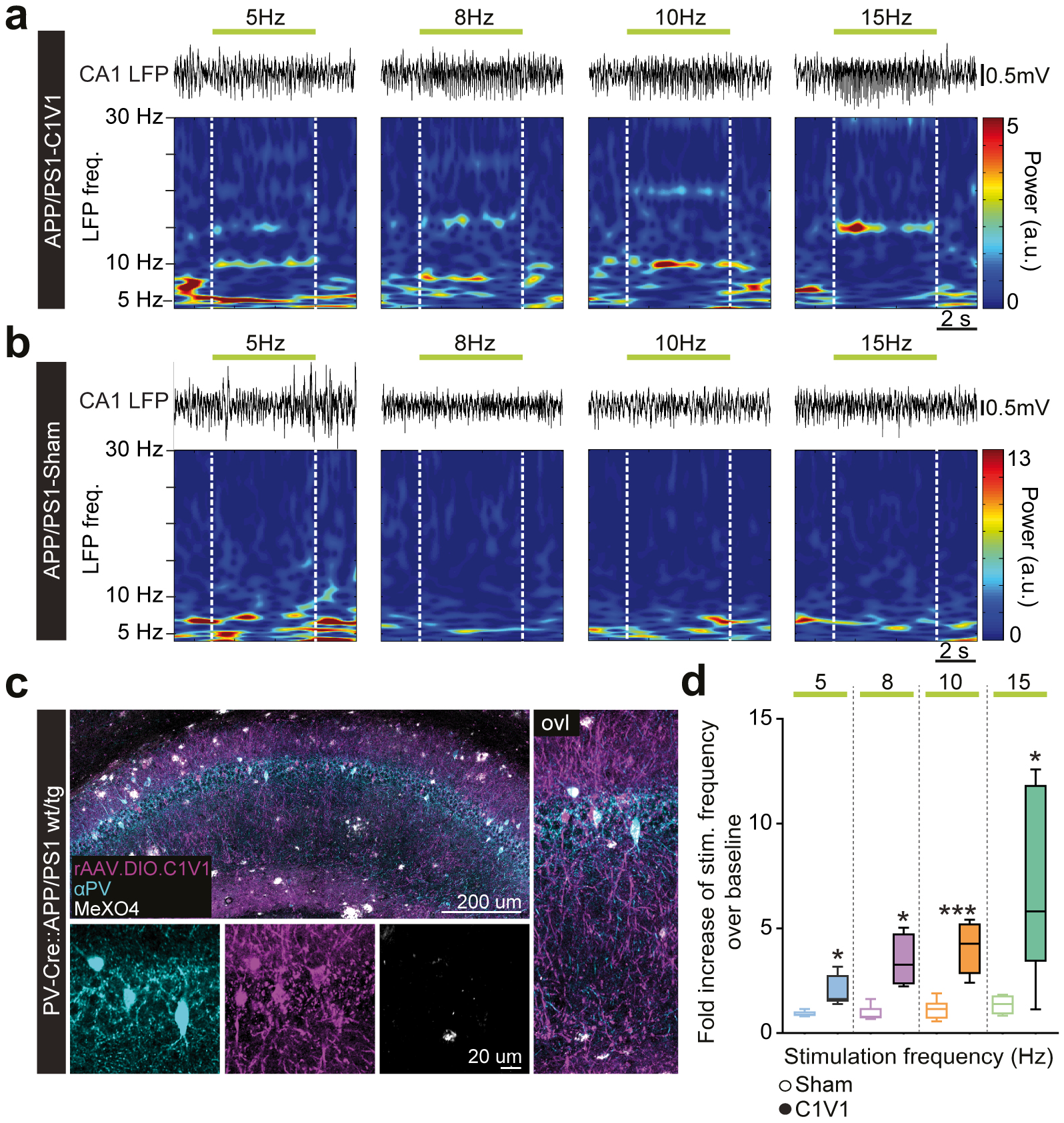
Optogenetic stimulation of PV^+^ interneurons drives LFP frequencies. This data was acquired in the mice employed for the arrhythmic stimulation (figure 3 **e** to **h**). **(a, b)** Exemplary spectrograms showing the effect of 5, 8, 10 and 15 Hz laser stimulation of the LFP of a C1V1-(a) and a Sham-injected (b) PV-Cre::APP/PS1 mouse. Each spectrogram corresponds to a single stimulation epoch. **(c)** Exemplary immunofluorescent staining showing colocalization of the rAAV.DIO.C1V1 with the αPV antibody. Aβ plaques are stained by the MeXO4 dye. **(c)** Fold increase of stimulation frequency over baseline. Sham: n= 5-6 mice per frequency; C1V1: n= 4-5 mice per frequency. ^*^p<0.05, unpaired Student’s t-tests with Welch’s correction; ^***^p<0.01, unpaired Student’s t-test. The dashed line represents the baseline to which the fold increase was normalized. Whisker plots represent median, 25^th^ and 75^th^ percentiles. Individual mouse values result from the average of 2-6 stimulation protocols.

**Supplementary figure 5.**
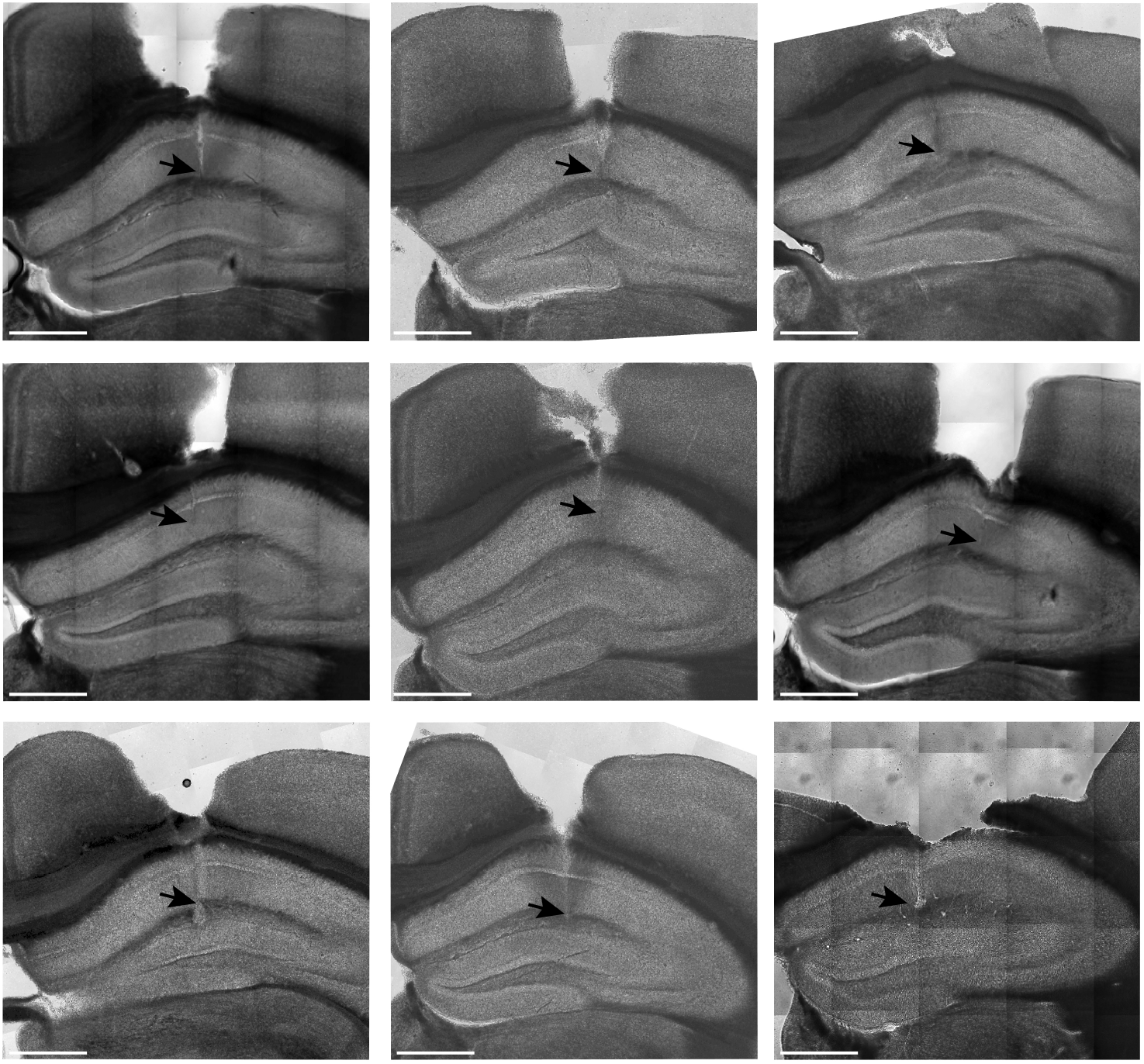
Examples of electrode and light fiber placement. Brightfield confocal images showing position of the electrode tip in the stratum radiatum of the hippocampal CA1 area of 9 exemplary mice (as indicated by the arrowheads). Only one of the two hemispheres is shown. Scale bars: 1mm in all panels.

## Materials and methods

### Animals

Experiments were performed on male and female PV-Cre::APP/PS1 mice. These were generated by crossbreeding B6.129P2-Pvalb^tm1(cre)Arbr^/J mice (named PV-Cre, Stock number: 008069, The Jacksons Laboratory) to B6.Cg-Tg(APPswe,PSEN1dE9)85Dbo/Mmjax mice (named APP/PS1, MMRRC stock number 034832, The Jacksons Laboratory). Mice were housed with their littermates under specific pathogen free conditions and were given food and water *ad libitum*. All experiments were conducted during the light phase of the 12 h dark/light cycle (lights on at 6:00 am, lights off at 6:00 pm) in the same room, lit by a normal neon lamp. The experimental procedures were in accordance with guidelines established by the DZNE and were approved by the government of North-Rhine-Westphalia. At the end of the experimental procedures, wild type mice were aged 14 ±3.1 months and transgenic mice 15 ±3.5 months (mean ±SD) for experiments shown in Figures 1, 2 and 3 e-h. For the second cohort of mice, we employed only APP/PS1 transgenic animals (Figure 3 h), and average age at the end of experimental procedures was 14 ±2.2 months (mean ±SD).

### Surgical procedures

#### Stereotactic viral injections

Mice were anesthetized with a mixture of ketamine/xylazine (0.13/0.01 mg/g body weight; Bela Pharm GmbH&Co. KG, Vechta and Rompun 2%, Bayer Vital GmbH, Leverkusen, DE) injected intraperitoneally. Additionally, mice received temgesic as analgesic (0.05 mg/kg; RB Pharmaceuticals Limited, Mumbai, IN), and dexamethasone as anti-inflammatory drug (0.2 mg/kg; Sigma Aldrich, Taufkirchen, DE), all injected subcutaneously prior to the surgery. A layer of eye-ointment (Bepanthen, Bayer Vital GmbH, Leverkusen) was applied on the eyes of the mice to prevent de-hydration. Upon absence of toe-pinch reflex, a small skin incision was made to expose the skull. Two small holes (~0.5 mm diameter) were opened through the skull using a dental drill. *AAV2-Ef1a-DIO C1V1 (t/t)-TS-mCherry* and *AAV1-CAG-FLEX-tdTomato* were bilaterally injected in the hippocampi of different groups of wild type and transgenic PV-Cre:APP/PS1 mice. 1 μl of virus was injected in each hemisphere at a speed of 100 nl/min, above the CA1 area (AP: −1.8 mm from bregma, ML: +-1.5 mm from midline and DV: +1.1 mm from brain surface). Injections were performed using a Hamilton syringe with a 34G needle (World Precision Instruments, Berlin, DE). Injection volume and speed were regulated using an UltraMicroPump (World Precision Instruments, Berlin, DE), and a micromanipulator (Luigs and Neumann, Ratingen, DE) was employed for stereotactic motion control. The needle was kept in the brain for 10 minutes prior to retraction. Finally, the skin was sutured with surgical wire (Vicryl, Ethicon, Norderstedt, DE). Mice were provided with analgesic (0.05 mg/kg) during three consecutive days following the surgery. Double fiber optic cannulas (light guides) and electrodes were implanted earliest 1 week and latest 3 weeks after stereotactic AAV-injections.

#### Optic ferrules and LFP electrodes implantation

Mice were anesthetized and provided with analgesic, antibiotic and anti-inflammatory as described above. Double fiber optic cannulas (B284-3015-2.5, TFC_300/370 – 0.22-2.5 mm, TSM3_FLT, Doric lenses, Quebec, CA) were implanted above the *stratum oriens* of CA1. Electrodes were custom made by soldering a tungsten wire (bare wire: 0.075 mm, coating thickness quadruple PTFE: 0.018 mm, 99.98% purity, ADVENT Research Materials ltd., Oxford, EN) into a gold pin (Conrad, DE). Prior to implantation, electrodes were glued to the optic cannulas using a drop of dental cement (Cyano Veneer Pulver A2) mixed to the special glue (Cyano Veneer, Fast-retarder, Hager & Werken GmbH & Co. KG, Duisburg, DE). Electrodes were cut 250 μm longer than the optic cannulas to target the *stratum radiatum* of CA1. This ferrule-electrode arrangement was stereotactically implanted at the same AP and ML coordinates as the injection sites but at a depth of 900 μm from brain surface (from the optic fiber’s tips). Reference and ground electrodes were additionally implanted in the cerebellum through small holes drilled into the cerebellar skull bone. Light guides and electrodes were fixed to the skull using a mixture of dental cement (Cyano Veneer Pulver A2) and special glue (Cyano Veneer, Fast-retarder, Hager & Werken GmbH & Co. KG, Duisburg, DE). Mice were allowed at least one week of post-surgical recovery before experimental procedures began.

### Electrophysiological recordings in brain slices

#### Slice preparation

Hippocampal coronal slices (300 μm thick) from wild type PV-Cre::APP/PS1 mice expressing *rAAV2-Ef1a-DIO C1V1 (t/t)- TS-mCherry* in the hippocampus were prepared with a Leica VT-1200S vibratome (Leica Microsystems, Wetzlar, Germany) in ice-cold sucrose solution containing (mM): 60 NaCl, 100 sucrose, 2.5 KCl, 1.25 NaH_2_PO_4_, 26 NaHCO_3_, 1 CaCl_2_, 5 MgCl_2_, 20 glucose, oxygenated with 95 % O_2_ and 5 % CO_2_. After recovery for at least 30 min at 35 °C slices were transferred into standard ACSF with the following composition (mM): 125 NaCl, 3 KCl, 1.25 NaH_2_PO_4_, 26 NaHCO_3_, 2.6 CaCl_2_, 1.3 MgCl_2_, 15 glucose) at room temperature.

#### Electrophysiological recordings with optogenetic stimulation

Electrophysiological recordings were obtained from neurons visualized by infrared DIC and fluorescence microscopy for mCherry identification (SliceScope with BX-RFA, Scientifica, East 25 Sussex, UK, Olympus, Hamburg, Germany). After confirmation of mCherry positive neuron in hippocampal CA1 area patch-clamp recordings were performed. Whole cell patch clamp recordings of either PV+ interneurons or pyramidal neurons in hippocampus were performed using a ELC-03XS amplifier (npi electronic, Tamm, Germany) and digitalized at 50 kHz sampling rates using an ITC-18 interface board (HEKA) controlled by self-made acquisition software based on Igor Pro 6.3 (WaveMetrics, Portland, USA). PV^+^ interneurons and pyramidal neurons were identified by mCherry expression and by the morphology of the cells, respectively. The cell types were also confirmed by the firing properties afterwards. The recording pipettes with a resistance of 4-6 MΩ were filled with standard intracellular solution containing (mM): 140 K-gluconate, 7 KCl, 5 HEPES-acid, 0.5 MgCl_2_, 5 phosphocreatine, 0.16 EGTA. All recordings were performed at 34 °C without correction for liquid junction potentials. Immediately after establishing the whole-cell configuration, the resting membrane potential (RMP) was measured. The membrane potential was adjusted to −60 mV by continuous current injection and a set of 500 ms hyper- and depolarizing stimuli (from –200 to +500 pA) was applied. Optogenetic stimulation was performed with a 561 nm continuous wave solid-state laser (OBIS 561 80 LS FP, Coherent, Santa Clara, USA) coupled to a light fiber. The light fiber tip was placed in a distance of ≤ 5 mm to the slice. Cells were stimulated with 8.5 Hz (15 ms pulse width) either without or with ≤30 ms random jitter of light for 5 s

#### Immunohistochemistry

At the end of the experimental procedures, mice were transcardially perfused using 1xPBS and their brains were fixed overnight in 4% paraformaldehyde (PFA, Carl Roth GmbH + KG, Karlsruhe, DE). Afterwards, coronal slices were cut using a vibratome (Leica VT 1200S, Leica Byosystems Nussloch GmbH, Heidelberg, DE) at a thickness of 100 μm. PV^+^ interneurons were stained using an α-PV^+^ primary antibody (1:1000, PV-25, Swant, Marly, CH) diluted in the permeabilizing solution: 2% bovine serum albumin (2% BSA, Carl Roth GmbH + Co. KG, Karlsruhe, DE), 2% Triton X100 in 1xPBS (AppliChem GmbH, Darmstadt, DE) and 10% normal goat serum in 1x PBS (10% NGS, Thermo Fisher Scientific, Darmstadt, DE). Slices were incubated overnight with the primary antibody solution at room temperature on a shaker (300 rpm). On the next day, the solution was removed and the slices were washed in 1xPBS. A rabbit anti-mouse-Alexa488 antibody, diluted in 3% BSA/PBS, was used as secondary antibody (1:400, rabbit α-mouse, Invitrogen, Darmstadt, DE) for 2.5 hours at room temperature (300 rpm). Upon removal of the secondary antibody solution, slices were washed with 1% TritonX100 in 1x PBS for 10 minutes and finally with 1x PBS. Slices were mounted using fluorescent mounting medium (Dako, Deutschland GmbH, Hamburg, DE). Confocal microscopy pictures were acquired using a LSM700 system (Carl Zeiss Microimaging GmbH, Jena, DE).

### Behavioral procedures

#### LFP recordings in the open field

LFPs were recorded while mice explored a squared open field (50×50 cm). Mice were let freely move in the arena for a maximum of 60 minutes and were videotaped for offline analyses. LFPs were sampled during the whole session. Mice were recorded until a minimum of 2 sweeps reaching a threshold activity criterion was reached (total travelled distance >100 cm). A custom-made faraday cage and a white curtain surrounded the open field, and no specific cues were added. The arena was wiped with 70% ethanol and water after every mouse.

#### Optogenetic stimulation in the open field

These experiments were performed in the same open field arena, room, and with the same conditions as mentioned above (open field). To verify that light stimulation of PV+ interneurons could drive theta frequencies *in vivo*, we stimulated these interneurons at 5, 8, 10 and 15 Hz with a 561 nm wavelength laser (OBIS 561 LS – laser system, Coherent, Santa Clara, USA) while the mice freely explored the environment. Laser power and pulse duration were kept constant at 20 mW and 15 ms, respectively. Power at the end of each optic fiber was 9 ±1 mW. Light was bilaterally delivered to the brain using a rotating connector that served as a laser beam splitter. Light was equally distributed into two fiber optic patch cords (P54997-04, CM3-SMC, core: 300 um, NA:022, Doric lenses, Quebec, CA), which were in turn connected to the optic fiber adaptors on the intracranially implanted ferrule. After every mouse, the arena was wiped with 70% ethanol and water.

#### Novel object recognition

Mice performed a novel object recognition (NOR) test in a squared arena (50×50 cm) as described above (open field). During the sample phase, mice explored the arena, which contained two identical objects (two flasks filled with brown soil) placed in the center. The sample phase consisted of 3 exposures to the arena (6 minutes each), interspaced by 15 minutes. After 24 hours, one of the two familiar objects was replaced by a novel one (a metallic cylinder), and the mice were placed back in the arena for 10 minutes. The site of entrance into the arena and the position of the novel object were pseudo-randomized between groups. During both sample and test phases, LFPs were recorded and a feedback optogenetic stimulation protocol was applied (*see stimulation protocols*). LFPs were acquired from only one of the two recording electrodes. After every mouse, the arena and the objects were wiped with 70% ethanol and water.

### LFP recordings and stimulation protocols

#### Setup

All electrodes were connected by means of custom assembled 2 m long insulated cable (single conductor wire LifY, 0.05 mm^2^ and 0.8 mm insulation, Conrad, DE) ending in a pin-socket on one end and connected to the headstage on the other. LFPs were sampled at 25 or 10 kHz using an amplifier system and two portable headstages (NPI electronic GmbH, Tamm, DE). The signal was low-pass filtered at 700 Hz and digitalized with an ITC-18 interface board (HEKA, New York, USA). The IgorPro software v. 6.22A (Wavemetrics, Portland, USA) was used for data collection and control of the laser stimulation. LFP data was downsampled to 1 kHz for offline analysis.

#### Closed-loop stimulation protocol

##### Rhythmic stimulation

Using a custom designed procedure in IgorPro, a closed-loop stimulation was established to modulate theta frequencies in the local field potential: In intervals of 500 ms a Fourier transform of the past 4 seconds of LFP recording was calculated using a fast Fourier transformation. A clear peak in the theta band (4-12 Hz) was defined as a ratio of the peak theta amplitude to the average theta amplitude exceeding 4. Whenever this criterion was fulfilled, a stimulation interval at the detected peak frequency plus 0.5 Hz with 15 ms pulse duration was initiated and protracted for 5.5 seconds. After each stimulation event, no new stimulation was initiated for at least 5 seconds to avoid detection of the induced activity.

##### Arrhythmic stimulation

The protocol described above was modified with a random jitter that followed a uniform distribution in the interval [-30 ms, +30 ms] to shift the timing of light pulses. This caused the randomization of the pulses across the stimulation interval. Every stimulation interval produced a different randomized set of pulses. However, on average, the stimulation frequency was the detected theta frequency plus 0.5 Hz as during the rhythmic stimulation.

#### Optogenetic stimulation in the open field

Stimulation protocols were designed in IgorPro. A 20 second protocol was employed to test whether stimulation at 5, 8, 10 or 15 Hz would increase the amplitude of the respective stimulation frequency in the LFP. The protocol consisted of 5 seconds of baseline followed by 5 seconds of stimulation and further 10 seconds of recording in absence of stimulation. Laser power (20 mW) and light pulses (15 ms) were kept constant.

### Data analysis

LFP data was analyzed using custom-made MATLAB scripts (R2013B, Mathworks, Natick MA, USA) or with IgorPro v. 6.22A (Wavemetrics, Portland, USA). Only one of the two implanted electrodes was used for LFP recordings throughout all the behavioral experiments.

#### Analysis of whole-cell recordings

All whole cell patch clamp recordings were analyzed using Igor Pro 6.3. From first to tenth of IPSPs in pyramidal neurons were analyzed for IPSP peak amplitude and area. IPSPs area was calculated for 55 ms from onset of light stimulation.

#### Baseline recordings

To detect differences in theta parameters between wild type and transgenic animals, values were extracted from spectrograms produced with the wavelet method in MATLAB, using the FieldTrip toolbox^27^. Theta parameters (peak theta frequency, amplitude and duration of theta phases) were calculated for each recording sweep (3 minute per sweep). Only sweeps where the mice were actively exploring were included in the analysis (total travelled distance >100 cm). Intervals of theta oscillations were defined as epochs when theta amplitude (5-12 Hz) increased by a ratio of 6 (arbitrarily set) over the delta band (4-5 Hz). The peak theta frequency was first averaged over sweeps for every mouse (minimum 2 sweeps, maximum 6). Subsequently, these values were averaged for each genotype. Travelled distance, mean velocity and time spent in the center were automatically analyzed in EthoVision XT (Noldus, Wageningen, NL). Center of the arena was defined as the squared portion of the open field being 5 cm distant from every wall of the arena.

#### Optogenetic stimulation in the open field

Fourier transformations of baseline and stimulation intervals were computed with IgorPro. The peak amplitude of the frequency of interest (5, 8, 10 or 15 Hz) was normalized to baseline intervals (5 seconds prior to stimulation), and fold increase in the peak amplitude of the frequency of interest over baseline during stimulation intervals was calculated. Four to six stimulation protocols were applied per mouse, and the fold increase of the frequency of interest over baseline was averaged first per mouse, then per treatment group.

#### Novel object recognition

Travelled distance, total exploration time, mean velocity and time spent at objects were measured in EthoVision XT (Noldus, Wageningen, NL). An area of ca. 3 cm was drawn around both familiar and novel objects, and nose point entry into that arena was considered as object exploration. The *discrimination index* (DI) was calculated as the percentage of time spent exploring the novel object over the total exploration time: (novel object exploration time / total exploration time)^*^100%. The behavior of the mice was analyzed for 5 minutes.

#### Statistical analyses

Statistical tests were carried out with GraphPad Prism 7 (GraphPad Software Inc., California, USA). Two-tailed unpaired Student’s t-tests were used to compare theta parameters (Fig. 1c and d;, Supplementary fig. 1a) and movement parameters (Figure 1e, f). A two-tailed Mann-Whitney test was used to compare not normally distributed theta power values (theta/delta ratio; Supplementary figure 1b). Two-tailed unpaired Student’s t-tests were used to compare effect of light stimulation at each frequency (5, 8, 10, 15 Hz) between Sham and C1V1 expressing mice (Figure 2d and Supplementary figure 4d). Differences in discrimination indices during the NOR test (Figure 3g) and locomotion parameters (Supplementary figure 2b) were compared using a two-way ANOVA followed by Holm-Sidak’s multiple comparison correction. Two-tailed unpaired Student’s t-tests were used to compare DIs and travelled distances of mice stimulated with the arrhythmic feedback stimulation protocol (Fig. 3h and Supplementary figure 2d). A two-tailed unpaired Student’s t-test was used to compare regression slopes (Supplementary figure 3e and k). All data was tested for normal distribution using the D’Agostino & Pearson normality test. When the number of values was too small to carry out a normal distribution test, normal distribution was assumed.

